# Cortical areas for self-recognition in the monkey brain

**DOI:** 10.64898/2026.11.19.767782

**Authors:** Tianyi Jiang, Shuaikang Xie, Liqing Tao, Haoyue Qian, Junfeng Huang, Liangtang Chang, Pengmin Qin, Zhifeng Liang, Yan-Yu Zhang, Jun Gao, Le Chang, Neng Gong

## Abstract

The sense of self represents a core feature of human cognition. Whether and how the monkey brain supports mirror self-recognition (MSR) remains unknown. Here, we show that monkeys acquired MSR following visual–proprioceptive association training, which involved stepwise learning to touch light spots—first directly visible and then, on an adjacent board, visible only in the mirror—and ultimately resulted in spontaneous mirror-guided face/body mark-touching and self-directed behaviors. In three MSR-capable monkeys, fMRI revealed that, relative to familiar and unfamiliar monkey faces, viewing the self-face elicited selective activation in the secondary somatosensory and insular cortices (SII-insula), known sites for multisensory integration. Longitudinal imaging throughout the training revealed that such selective SII-insula activation emerged only when the monkeys exhibited MSR. Dynamic causal modeling further showed self-face-specific modulation of the SII-insula node at this stage. Thus, the SII-insula constitutes a fundamental neural unit underlying self-recognition in the primate brain.

## Introduction

The neural mechanism by which the brain generates this sense of self remains largely elusive. In recent years, a growing body of research has sought to explore the nature of the “self”, originally a philosophical topic, through methods of cognitive neuroscience^1–3^. In particular, bodily self-consciousness (BSC), as the most foundational level of self, can be experimentally examined by manipulating multisensory integration^4,5^. For example, paradigms such as the rubber hand illusion and out-of-body experiences demonstrate that repetitive and synchronous visual and tactile stimulation can induce illusory percepts of body ownership^6,7^, self-identification^8,9^, and self-location^10,11^. Specific patterns of brain activity associated with subjective bodily self-illusions have been identified in various brain regions, including the premotor cortex (PMC), intraparietal sulcus (IPS), and temporoparietal junction (TPJ), where exteroceptive bodily signals are integrated^8,9,12^, as well as the insula that is engaged in processing interoceptive signals^13,14^. The coupling of internal and external bodily information thus appears critical for the emergence of a coherent bodily self^2,15,16^. In humans, distinct brain areas may be involved in self-consciousness at different levels^17–19^, but it remains unclear whether a core self-processing network exists in the brain. Research employing animal models may offer key insights into the neural substrate of self-consciousness and its evolutionary origin.

Whether animals possess a sense of self comparable to that of humans has long been debated^20,21^. In previous studies, macaque monkeys were shown to acquire mirror self-recognition (MSR) ability through visual-somatosensory/proprioceptive association training^22,23^. These trained monkeys passed the mirror mark test and spontaneously displayed mirror-induced self-directed behaviors—an ability that typically emerges in human infants at around 18 months of age^24^—thereby providing a non-human primate model for studying the neural mechanisms underlying self-recognition^25–27^. A useful approach is to examine changes in brain activity during the entire training process for the acquisition of MSR ability in monkeys. This would enable the longitudinal observation of the emergence of self-recognition in the same individual, an approach difficult to implement in human studies. Furthermore, comparison of brain regions activated during self-recognition in MSR-capable monkeys with those found during self-processing in humans may shed light on the evolution of self-consciousness. In this study, we employed functional magnetic resonance imaging (fMRI) in awake monkeys to compare cortical activation elicited by exposures to visual stimuli of self, familiar, or unfamiliar monkey faces throughout the MSR training process. We identified secondary somatosensory and insular cortices (SII-insula), cortical regions known to be critical for multisensory integration, as the primary brain areas for self-recognition in the primate brain.

## Results

### Behavioral stages of MSR training in monkeys

To investigate longitudinally the neural mechanism underlying self-recognition in macaque monkeys, we first characterized the behavioral progression throughout the MSR training and divided the process into distinct stages (Figure 1A; Video S1). In the first stage, monkeys were trained to touch a visible red light spot projected on a board, a simple skill typically acquired within 1–2 days. In the second stage, the monkeys underwent visual–proprioceptive association training^23^ over several weeks and learned to use the mirror image to locate and touch light spots on a board adjacent to the monkey’s head. Notably, after mastering this skill (the end of second stage), the monkeys were incapable of touching light spots projected directly onto their own faces or bodies via viewing the mirror. This suggests that MSR ability involves more than simple spatial mapping of the real light spot via its mirror image and requires instead higher-level cognitive processing. In the third stage, over the subsequent weeks under the same visual-proprioceptive training as in the second stage, the monkeys began to spontaneously generalize the spot-touching behavior to their own faces and bodies. A full generalization of spot-touching to the whole body emerged spontaneously after a further period of several weeks. Afterwards, the monkeys exhibited typical spontaneous MSR behaviors in their home cages (Figure 1B; Video S1), including passing the mirror mark test and displaying spontaneous self-directed behaviors in front of the mirror^22,23^, marking a critical breakthrough in their bodily self-consciousness. In this study, three monkeys underwent visual–proprioceptive association training and successfully demonstrated MSR ability after 6–8 weeks of training. Their behavioral progression during training displayed clear stage-dependent characteristics (Figure 1C).

**Figure 1.**
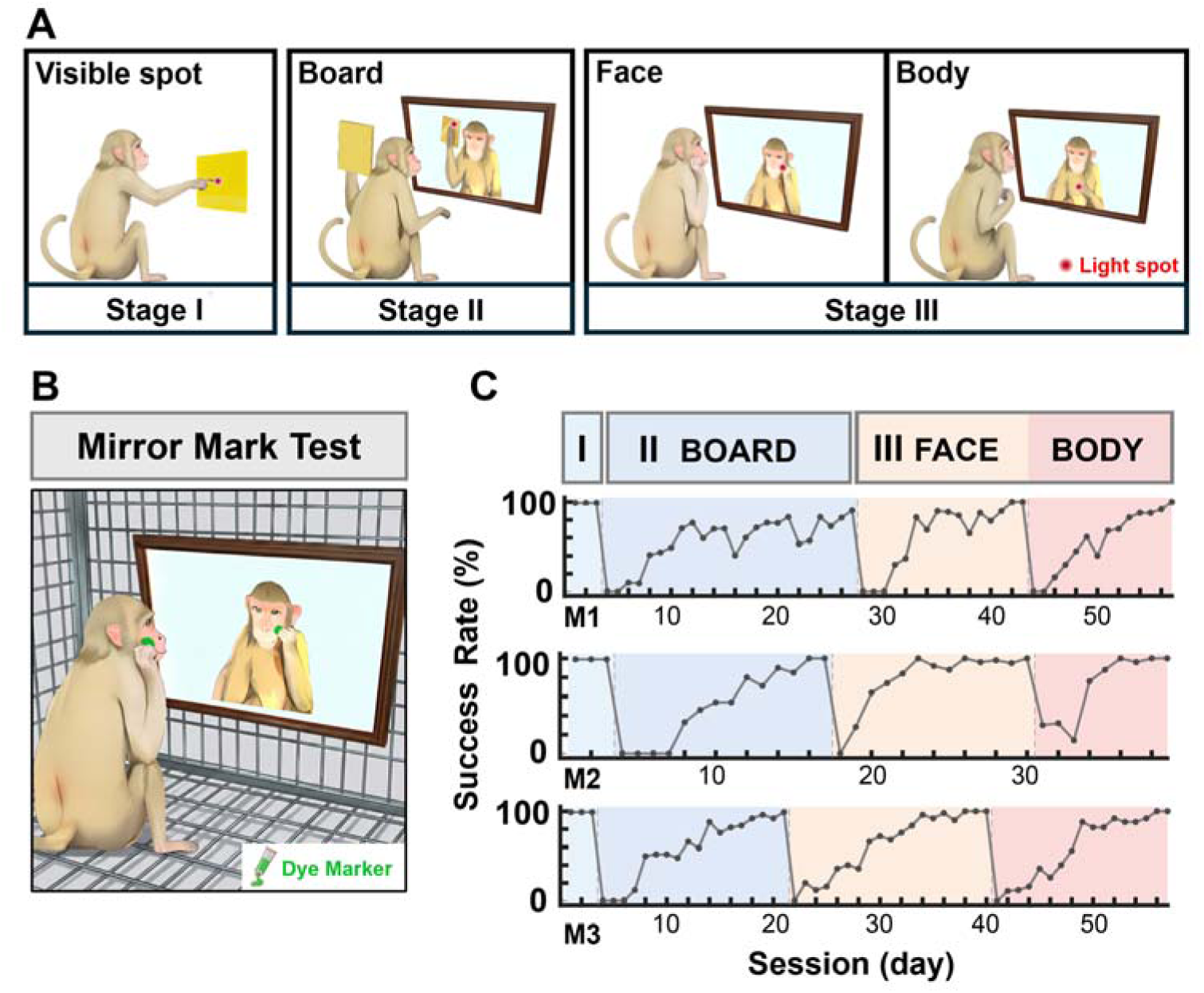
Behavioral stages of MSR training in monkeys. **(A)** Schematic of MSR training procedure. Stage I: Touching visible red light spots. Stage II: Touching light spots on the adjacent board in front of a mirror. Stage III: Touching light spots on their own faces and bodies in front of a mirror. **(B)** Illustration of the mirror mark test for the trained monkey in the home cage. **(C)** Behavioral performance across three stages of MSR training for the three monkeys used in this study.

### Selective cortical activation elicited by self-face in monkeys with MSR

Next we used fMRI to investigate the cortical regions activated throughout MSR training. We surmise that the monkeys’ perception of their own faces undergoes transitions from recognizing an unfamiliar to a familiar face and finally to self-face. For the fMRI study, we thus exposed the awake monkeys in the scanner to visual stimuli of static self-face (SELF) images as well as non-self monkey face images including visually familiar (VF) and non-familiar (NF) faces. The VF face stimuli were used to control the effects of facial familiarity during training and were obtained for each monkey as follows (Figure 2A). Alongside daily mirror training sessions, each monkey was also exposed to another monkey for an equivalent duration instead of the mirror, ensuring a comparable degree of visual familiarity between the self-face and this VF face of another monkey. Using these SELF, VF, NF and scrambled (SC) face images as visual stimuli, we performed awake-monkey fMRI on the three trained monkeys across all three training stages (Figure 2B). For each stage, once a sufficient number of fMRI runs (> 30 runs) were acquired, the monkeys proceeded to the next training stage; this procedure continued until they successfully passed the mirror mark test.

**Figure 2.**
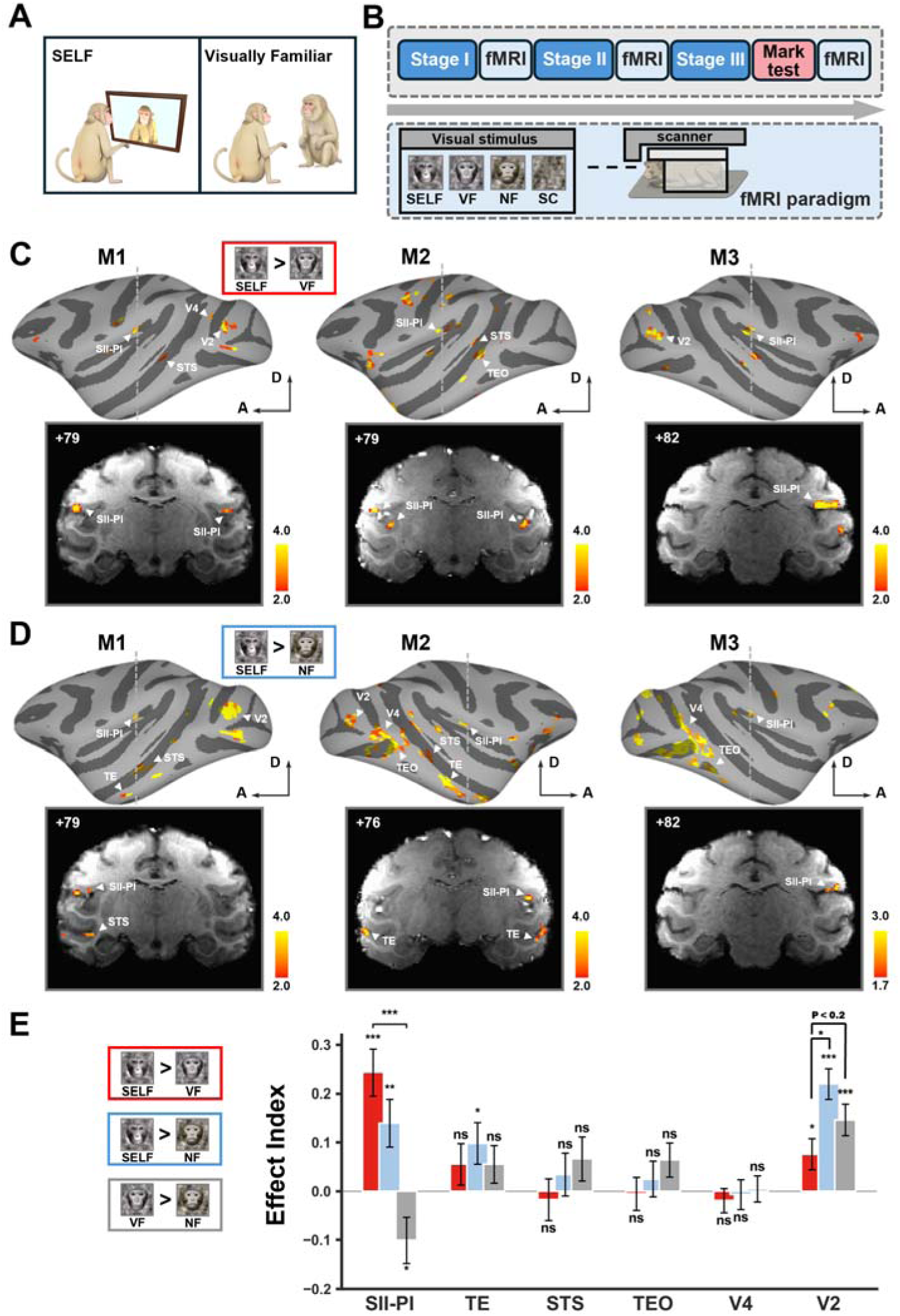
Cortical activations by self-face in monkeys with MSR. **(A)** Illustration of self-face (SELF) and visually familiar face (VF) stimuli. The VF face stimulus was derived from a monkey with a familiar-face condition (see supplementary materials). **(B)** Schematic of the fMRI scanning across all three training stages for all three monkeys in awake states. A display screen placed in front of the animal presented visual stimuli, including SELF, VF, unfamiliar (non-familiar, NF) faces, and face-matched scrambled (SC) stimuli. **(C)** Individual-level voxel-wise analyses contrasting SELF with VF after monkeys passed the mark test. (TOP) Inflated hemispheres showed the regions that responded significantly more to SELF than to VF using a Linear Mixed-Effects (LME) model for each monkey (M1, M2, M3) (uncorrected *P* < 0.01). The color bar denotes t-statistic values. (BOTTOM) Results are overlaid on coronal slices illustrated the significant position of the SII/PI complex in the SELF versus VF contrast. **(D)** Same as (C) but for SELF versus NF (uncorrected *P* < 0.01 for M1 and M2, uncorrected *P* < 0.02 for M3). **(E)** Contrast effect index for SELF versus VF, SELF versus NF, and VF versus NF of each region of interest (ROI) in monkeys with MSR ability. The effect index for face X versus face Y was calculated as [β(X) − β(Y)] / (abs[β(X)] + abs[β(Y)]), based on pooled data from all three monkeys. Error bars represent the standard error of the mean (SEM) across runs. \**P* < 0.05, \*\**P* < 0.01, \*\*\**P* < 0.001, ns: not significant. SII/PI, the secondary somatosensory cortex and posterior insula complex; V2, visual area 2; V4, visual area 4; TE, anterior inferotemporal cortex; TEO, posterior inferotemporal cortex; STS, superior temporal sulcus.

We first analyzed the fMRI data obtained after the monkeys had passed the mark test, the stage at which the self-face was supposed to elicit the sense of self in the monkeys. Comparing the activation elicited by SELF versus VF across the three monkeys revealed significantly higher activation in two sets of cortical regions: one encompassing the secondary somatosensory cortex (SII) and the posterior insula (PI), and the other comprising visual areas such as the visual area 2 (V2) in the visual cortex and face-selective patches in the temporal cortex (Figure 2C). Because of the spatial proximity of SII and PI, along with the limited spatial resolution of the fMRI data, we combined these two areas into a single complex, termed “SII-PI”, for subsequent analyses. Notably, higher activation in SII-PI in the SELF vs. VF contrast was highly consistent across individuals, with similar stereotactic coordinates in both hemispheres (Figure 2C; Table S1). Moreover, higher activation of SII-PI was also observed when comparing cortical activation elicited by SELF versus NF faces in all three monkeys (Figure 2D; Table S1), whereas activation in visual areas was substantially stronger in the SELF vs. NF contrast than in the SELF vs. VF contrast (Figures 2C and 2D). Higher activation of visual areas was also observed for VF vs. NF contrast (Figure S1), supporting the involvement of visual areas in processing facial familiarity^28,29^. Quantitative ROI-based analyses of the effect index (see Methods) for the contrasts of SELF vs. VF, SELF vs. NF, and VF vs. NF showed that significant effects were largely confined to the SII-PI and V2 (Figure 2E). Specifically, the SII-PI exhibited significantly higher activation mainly when the self-face was compared with either of the other two face types, whereas V2 activation was higher when SELF and VF were contrasted with NF faces (Figure 2E). Thus, in MSR-capable monkeys, self-face specifically enhanced activation in the SII-insula, while V2 activation primarily reflected visual familiarity of faces.

### Longitudinal analysis of cortical activation across three behavioral stages

Given the finding that the activation of SII-insula and V2 was associated with the processing of self-face and face-familiarity, respectively, we predicted that SII-PI activation elicited by the self-face should emerge only when the monkey acquires the capacity for MSR, whereas V2 activation would exhibit a progressive increase over the course of training. This prediction was validated by our longitudinal cross-stage comparison of cortical activation elicited by SELF versus VF or NF faces (Figure 3A). Specifically, significantly higher SII-PI activation in SELF vs. VF and SELF vs. NF contrasts were observed exclusively at the third stage (Figure 3B). In contrast, activation in visual areas, especially in V2, increased gradually across the three stages (Figure 3B). Notably, during the initial training stage, activation levels in various visual areas were comparable for SELF and NF faces (Figure 3B). However, VF faces elicited significantly stronger activation than self-face stimuli in several visual areas, such as V2, visual area 4 (V4), superior temporal sulcus (STS) and posterior inferotemporal cortex (TEO) (Figure 3B). At the onset of training, the monkeys had no prior exposure to their own faces, whereas the familiar monkeys housed in the same colony were already visually familiar to the subjects. Thus, these first-stage observations in visual areas further support their involvement in mediating visual familiarity.

**Figure 3.**
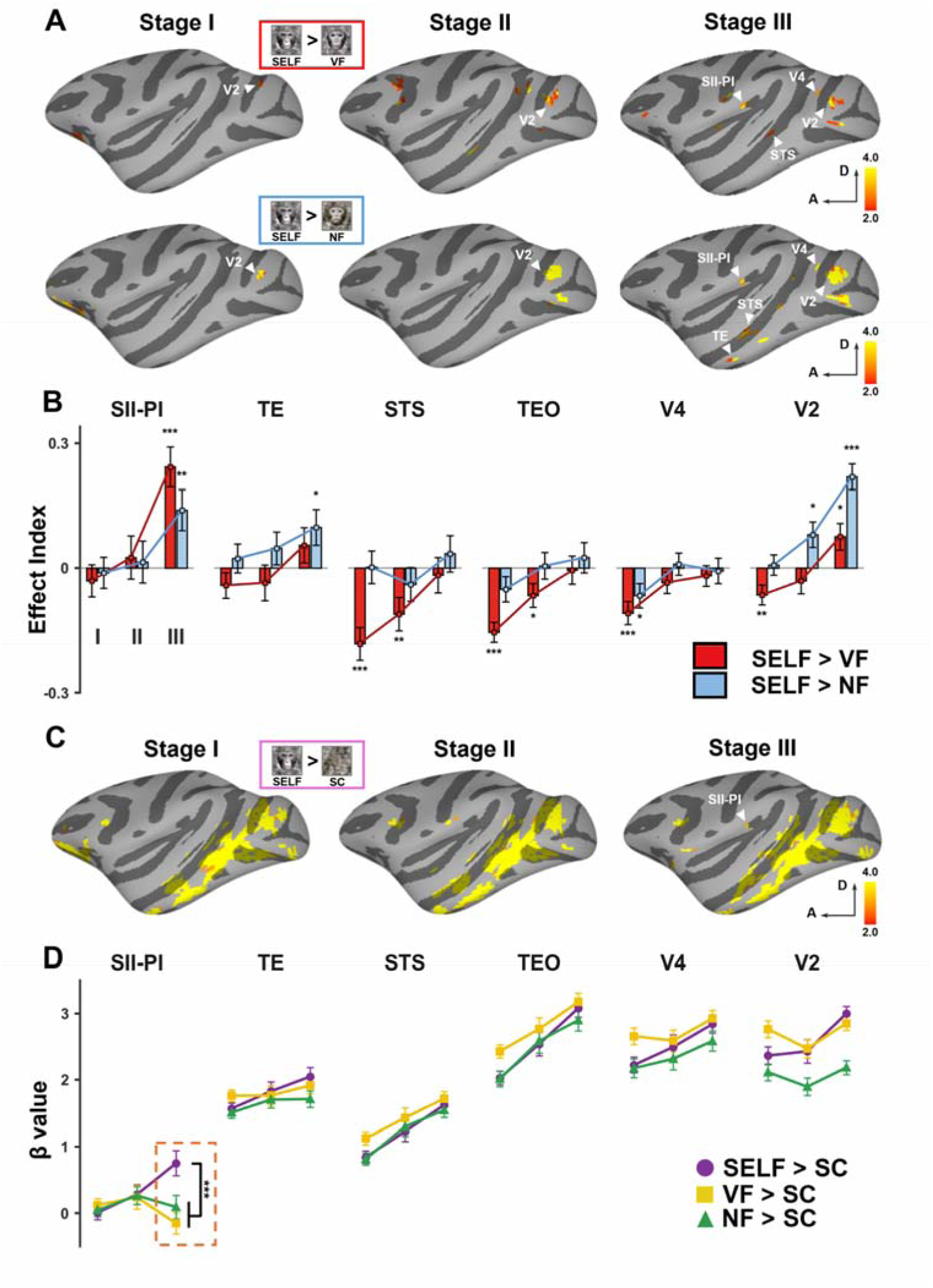
Cross-stage analysis of cortical activations during monkey MSR training. **(A)** Individual-level voxel-wise analysis for M1 across three stages. Inflated hemispheres showed the regions that responded significantly more to SELF than to VF/NF using an LME model (uncorrected *P* < 0.01). The color bar denotes t-statistic values. **(B)** Contrast effect index analyses for the selected ROIs across three stages for SELF versus VF, SELF versus NF, and VF versus NF. The effect index for face X versus face Y was calculated as [β(X) − β(Y)] / (abs[β(X)] + abs[β(Y)]), based on pooled data from all three monkeys. **(C)** Inflated hemispheres show regions with significantly greater responses to SELF than to SC for M1 across three stages, based on an LME model (uncorrected, *P* < 0.01). The color bar denotes t-statistic values. **(D)** Responses (β_FACE_ – β_SC_) to SELF, VF and NF stimuli in selected ROIs across three stages, based on pooled data from all three monkeys. Error bars represent the standard error across runs. \**P* < 0.05, \*\**P* < 0.01, \*\*\**P* < 0.001.

Finally, we performed an additional analysis of activation in the SII-PI and visual areas elicited by SELF, VF, and NF face stimuli, each contrasted with scrambled-face (SC) controls, across the three stages (Figure 3C). Relative to SC controls, face stimuli consistently evoked stronger and more spatially extensive activation across cortical areas, particularly in prefrontal and visual areas (Figure 3C). Notably, only in the third stage did SELF vs. SC contrast in SII-PI significantly exceeded the corresponding VF vs. SC and NF vs. SC contrasts (Figure 3D). In visual areas, all three face types exhibited similar changes in activation across the three stages when comparing with SC controls (Figure 3D). Together, these cross-stage cortical activation analyses further support the SII-insula as key brain areas involved in self-recognition in monkeys.

### Dynamic causal modeling across three behavioral stages

To further investigate how the SII-PI and V2 regions contributed to self and familiarity processing at the network level, and how effective connectivity was modulated by SELF, VF, and NF face stimuli, we performed dynamic causal modeling (DCM) analyses on fMRI data from the three behavioral stages (Figure 4A-4C; Figure S3). Specifically, the SII-PI complex was treated as a single node, the three face types served as driving inputs to V1, and a three-node model was constructed with bidirectional connections between V1 and V2 and between V1 and SII-PI, together with the self-connections of all three nodes, with the three face types serving as modulatory conditions for all connections. As shown in Figure 4A, in the group-level analysis across all runs of the three monkeys that had acquired MSR ability, the SELF and VF faces exhibited significant negative modulatory effects on the feedback connection from V2 to V1 (SELF modulation: −1.01; VF modulation: −0.93) relative to the NF face, with strong evidence (i.e., posterior probability > 0.85), demonstrating that functional connectivity from V2 to V1 was more strongly modulated by visual familiarity. In comparison, the SELF face, relative to the other two non-self faces, showed a significant negative modulatory effect on the feedback connection from SII-PI to V1 (modulation: −0.80), as well as a positive modulatory effect on the self-connection within SII-PI (modulation: 0.61), with strong evidence (i.e., posterior probability > 0.85). More importantly, this self-specific modulation of SII-PI-involved connections was absent in the first two training stages and emerged only after the monkeys acquired MSR ability (Figure 4B and 4C). Taken together, these results indicate enhanced intrinsic excitation in SII-PI, along with feedback inhibition of the primary visual area by SII-PI—mechanisms that may underlie self-recognition in the monkey brain.

**Figure 4.**
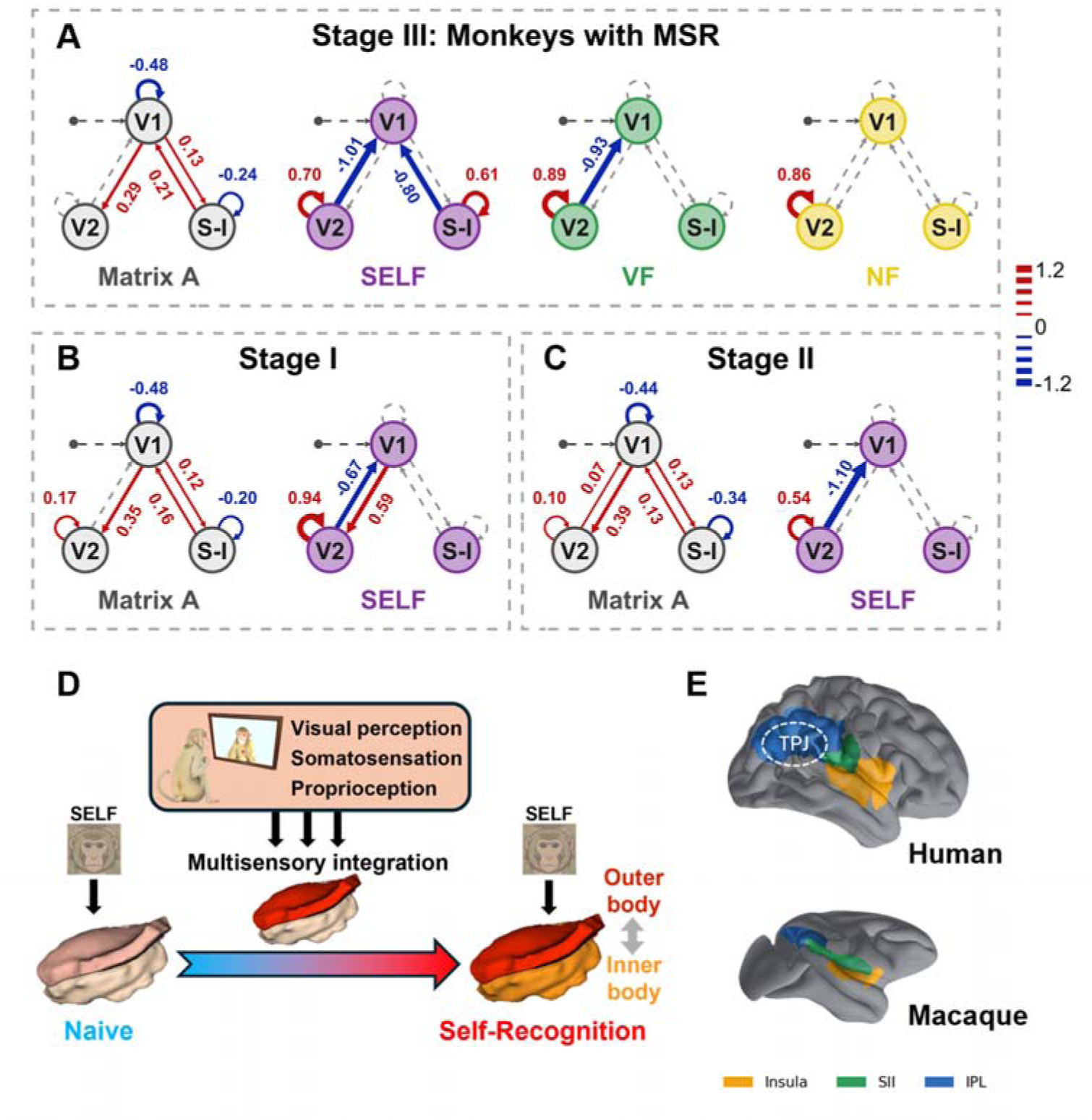
Stage-specific modulatory effects of SELF and hypotheses for self-processing. **(A–C)** Group-level dynamic causal modeling (DCM) results at Stage III (A; monkeys with MSR), Stage I (B), and Stage II (C). For each stage, the intrinsic connectivity of the three-node model (V1, V2, and S-I) is shown as “Matrix A”. Numbers are parameter estimates; colored arrows are shown for connections with posterior probability > 0.85, whereas gray dashed arrows indicate connections without significant modulation. The color bar denotes modulation strength (red, positive; blue, negative). S-I, SII-PI complex. **(D)** A working hypothesis for the function of SII-insula in the formation of bodily self-consciousness for monkeys undergoing MSR training. **(E)** Anatomical comparison between humans and macaques. Insula (orange), SII (green), and inferior parietal lobule (IPL, blue) are shown on the cortical surfaces, the dashed circle indicates the TPJ.

## Discussion

In this study, we found that the SII-PI region was specifically activated by self-face stimuli in monkeys capable of MSR. The SII is known to play a critical role in integrating external sensory signals and maintaining body representation^30–32^, whereas the insula is a key hub for interoception and is widely implicated in self-consciousness in human fMRI studies^16,33,34^. Notably, the SII-insula region in monkeys is also activated by pain stimuli^35,36^, which is likely to evoke potent self-experience. Furthermore, we did not detect self-face-specific activation in brain regions traditionally associated with conditioned learning, such as the prefrontal cortex, hippocampus, and striatum^37,38^, in support of the notion that the self-recognition behaviors demonstrated by the trained monkeys could not be attributed to conditioned learning^26,39^. Instead, it suggests that these monkeys genuinely experience a sense of self and possess a self-concept when viewing their own faces.

In monkeys with MSR, self-face also activated visual areas, but these responses appeared to be linked to face familiarity rather than being selectively associated with self-face recognition. In recent years, there is increasingly extensive and refined evidence supporting the role of visual areas in processing familiar faces^28,29^. In this study, visual area V2 was engaged in the processing of face familiarity at the final stage, which more likely reflects experience-dependent modulation of early visual encoding. We also found that in several higher visual areas, such as V4, TEO, and STS, the activation difference between the self-face and a familiar face became smaller as training progressed, suggesting that these brain areas are also involved in processing face familiarity at a higher level. However, it remained challenging to dissociate the contributions of visual familiarity and personal familiarity^28^, limiting our interpretation of familiarity processing in the current study. Nevertheless, our data did not provide any evidence that visual cortical regions or face-selective patches in the temporal cortex were specifically involved in self-recognition.

The DCM results further revealed the respective roles of the SII-PI and V2 in self and familiarity processing. At the stage when monkeys acquired MSR, both self and familiar faces exerted significant inhibitory modulatory effects on the V2–V1 feedback connection, whereas only self faces exerted an inhibitory modulatory effect on the (SII-PI)–V1 feedback connection and an enhancing effect on the self-connection of the SII-PI; these effects were absent in the first two stages of training. Drawing on these findings, we propose that MSR training with multisensory integration induced plastic changes in SII circuit functions that contribute to the self-recognition in monkeys (Figure 4D). In naïve monkeys, visual self-face stimuli alone are insufficient to activate SII for forming the sense of self. However, during the training, multisensory integration of visual feedback signals from the mirror, proprioceptive signals associated with the hand position, and somatosensory signals from whole-body touching led to functional modification of SII underlying the acquisition of MSR ability. During self-recognition, the activation of SII by self-face may subsequently engage the adjacent insular cortex, thereby coupling external sensory cues with internal bodily states. This cascade of neural activity ultimately underlies the emergence of self-recognition and the sense of self in monkeys (Figure 4D).

In line with the finding that the insular cortex is associated with the sense of self in humans^34,40^, we observed activation in the insula during self-face processing in monkeys with MSR ability. However, in these monkeys, we did not detect activation of TPJ, the key multisensory integration region associated with human BSC^9,12,41^. Instead, activation elicited by self-face in the monkey brain was identified in SII, a brain area known for its role in constructing a body map^30,31^. The SII also integrates various types of multimodal information, including visual, auditory, and vestibular inputs, and thus supports higher-order cognitive processes that require multisensory integration^32,42^. Indeed, human brain imaging studies have also indicated that SII may contribute to self-related cognitive functions such as self-awareness^7,43^, self-localization^44,45^, and self-recognition^32,46^. Iriki and colleagues have further proposed that SII functions as a site of information integration grounded in spatial body maps, and integrates diverse bodily sensations into a somatocentric holistic self, thus forming a cognitive representation of “self-in-the-world”^47^. Notably, SII neurons in MSR-capable monkeys responded to mirror image stimuli and displayed distinct activities in response to self vs. non-self stimuli^48^. The SII–PI region we identified in monkeys, together with the well-established TPJ and insular regions in humans, all lie along a cascading circuit comprising the insula, SII, and Inferior Parietal Lobule (IPL) (roughly equivalent to the TPJ in humans) (Figure 4E). We therefore propose that this cascading circuit constitutes a key circuit for interoceptive–exteroceptive integration and self-processing in primates. The information processing in monkey SII may represent a precursor of that in human TPJ, and this difference may reflect the evolutionary progression of self-related processing from monkeys to humans.

In macaque monkeys, a non-human primate with a brain structure similar to that of humans, we identified the SII-insula as key brain areas underlying self-recognition in monkeys. Our study demonstrates the utility of non-human primate models in probing the neural substrate and evolutionary origin of self-consciousness. This finding also underscores the importance of multisensory integration for the formation of self-consciousness and suggests that SII, the earliest cortical site integrating whole-body somatosensory inputs, serves as a fundamental unit enabling self-recognition.

## Acknowledgments

We thank Mu-ming Poo for helpful discussions and insightful comments on this manuscript, and Wenqiang Zhang for assistance in figure preparation. This work was supported by Brain Science and Brain-like Intelligence Technology – National Science and Technology Major Project (2021ZD0204200 and 2022ZD0204600), CAS “Strategic Priority Research Program” (XDB1010101), CAS Project for Young Scientists in Basic Research (YSBR-071).

## Author contributions

N.G., L.C. and J.G. conceived and designed the study. T.J., S.X., L.T., J.H. and L.C. performed monkey training and fMRI. T.J., S.X., L.T. and H.Q. analyzed data. T.J. and N.G. prepared the figures. N.G., T.J. and H.Q. wrote the manuscript. P.Q., Z.L. and Y.Z. provided critical suggestions on experimental design and data interpretation, and commented on the manuscript. All authors have read and approved the manuscript submission.

## Methods

### Subjects

We studied three adult rhesus monkeys (M1–M3, Macaca mulatta, weight 7–10 kg, 5–10 years old). A cranial implant of MR-compatible acrylic cements, anchored with MR-compatible ceramic screws, was placed in each subject following standard surgical, anesthetic, aseptic, and postoperative protocols. A custom-designed MR-compatible headpost was fixed to the implant. For fMRI scanning in awake monkeys, all monkeys underwent approximately four weeks of habituation training in an MRI-compatible monkey chair in the sphinx position, until they could remain calm and motionless for 2–3 hours continuously. Subsequently, they received around 6 weeks of fixation training in the chair. The fixation spot had a diameter of 0.2**°** of visual angle, and the allowed fixation window was a circular region with a diameter of 2.5**°**. Animal care and experimental procedures were approved by the Animal Care and Use Committee at the Institute of Neuroscience, Center for Excellence in Brain Science and Intelligence Technology, Chinese Academy of Sciences.

### Mirror self-recognition (MSR) training

Visual-proprioceptive association training was used for MSR in monkeys. This protocol followed the previous study^23^ and comprised three stages, with fMRI scans performed after each stage.

#### Stage 1: Touching visible light spots

Monkeys were trained to touch visible light spots while sitting in a custom-designed monkey chair. The laser was projected onto a board placed in front of them. Each successful touch resulted in an immediate fruit reward, reinforcing their behavior. The training continued until the monkey could reliably locate and touch light spots at various positions on the board.

#### Stage 2: Touching light spots on the adjacent board in front of a mirror

Monkeys learned to use mirror images to locate and touch light spots that were not directly visible. A mirror was placed in front of the monkey, and two boards were placed adjacent to the monkey’s head so that the monkey could see the board only in the mirror. The light spot initially appeared both directly and in the mirror, and then gradually moved to the adjacent board thus becoming visible only through the mirror. Upon successfully touching the light spot, the monkey received a reward. This training continued until the monkey could reliably locate and touch light spots at various positions on the adjacent board in front of a mirror.

#### Stage 3: Touching light spots on the monkey’s own face or body in front of a mirror

The light spot was projected to the monkey’s own face, and the monkey was allowed to spontaneously explore the position of the light spot. Upon successfully touching the light spot, the monkey received a reward. When the monkeys became proficient in touching spots on their own faces, the light spot was projected to other body parts including the chest, abdomen, and upper arms. Finally, the training context was shifted to the home cage where monkeys could freely touch spots on their entire body in front of a mirror.

### Mirror mark test

To assess MSR ability, a standard mirror mark test was conducted in the trained monkeys^49^. Odorless non-irritating skin-safe colored dyes or equivalent amount of water were randomly applied to several sites of the monkey’s face, such as the forehead, cheek, or ear. Monkeys were observed for 10–15 minutes without mirror, and then a mirror was reintroduced. Dye mark-touching and smelling, as well as self-directed behaviors in front of the mirror were considered as passing the test.

### MRI data acquisition

All MRI data were acquired in a 3T MRI scanner (Siemens Healthcare, Erlangen, Germany) while the monkeys were in sphinx position with a custom-built 8-channel surface coil. Each functional time series consisted of 156 gradient-echo echo-planar whole brain images [EPI; repetition time (TR) = 2000 ms; echo time (TE) = 24 ms; flip angle (FA) = 80°, matrix = 64×64; Field of view (FOV) = 96×96; voxel size = 1.5 mm×1.5 mm×2 mm; 27 horizontally oriented slices] acquired in interleaved order, and two times generalized auto calibrating partially parallel acquisitions (GRAPPA) acceleration. On each scanning day, a contrast agent (Molday ION, BioPAL, Worcester, Massachusetts, USA) was injected prior to scanning to improve the signal-to-noise ratio (SNR)^50^. High-resolution T1-weighted anatomical images were acquired using a magnetization-prepared rapid gradient echo (MPRAGE) sequence (TR = 2300 ms; TE = 2.7 ms; FA = 7°; matrix = 256×256; FOV = 128×128; voxel size = 0.5 mm×0.5 mm×0.5 mm, 256 slices).

### MRI stimuli and procedure

All monkeys completed passive image-viewing tasks during fMRI scanning, and four sets of stimuli were created for each subject to examine self-face recognition: [1] self-face (SELF). [2] visually familiar (VF) face obtained by a familiar-face condition. Specifically, alongside daily mirror training sessions, each monkey was exposed to another monkey for an equivalent duration, ensuring a comparable degree of visual familiarity. [3] unfamiliar (non-familiar, NF) face. [4] face-matched scrambled stimuli. To acquire images of face stimuli, we recorded 15–20 minutes of high-definition video of monkey faces and extracted 24 facial images captured from different views. Each image was then cropped using Adobe Photoshop to retain only the facial region—excluding all non-facial features such as ears—and standardized in terms of resolution, brightness, and size. The scrambled stimuli were created by phase-scrambling each face image: the phase component of the image Fourier transform was randomized and then inverse-transformed, preserving the power spectrum while disrupting fine spatial structure.

A typical fMRI block sequence was implemented for each monkey, where each block contained 24 images from one stimulus set presented in random order with a 0.5-s duration per image (each image repeated twice), and each scan consisted of 13 blocks lasting 312 s in total. For monkey M1, the scan included six face blocks (two SELF, two VF, and two NF blocks) and seven scrambled blocks, with the first block fixed as a scrambled block, followed by alternating face and scrambled blocks, and the order of the six face blocks was determined via a Latin square design with 24 possible sequences. For monkeys M2 and M3, the scan comprised five face blocks (SELF, VF and NF blocks were randomly assigned to one, two, and two blocks), one scrambled block, and seven grey baseline blocks (with the screen uniformly filled with grey), where the first block was fixed as a grey baseline block, followed by alternating face/scrambled and grey baseline blocks. The presentation order of the five face blocks and one scrambled block was also determined using a Latin square design with 24 possible sequences.

All stimuli (11°×11° visual angle) were presented on an MRI-compatible screen (1280×1024 pixels) positioned 92 cm from the monkey’s eyes, against a uniform gray background, with a central fixation spot measuring 0.2°×0.2° of visual angle. During the experiments, eye position and pupil diameter were monitored at 250 Hz using an MRI-compatible eye-tracking system (EyeLink 1000, SR Research, Mississauga, Ontario, Canada). A fluid reward was delivered after successful fixation, defined as maintaining gaze within a virtual central window of 2.5°×2.5° for a continuous duration of 3 seconds. Only fMRI scans in which the total fixation time exceeded 85% of the entire scan duration were included in subsequent analyses (Figure S2). Visual stimulation and reward were controlled by MATLAB scripts.

### Data analysis

Data analysis was performed using AFNI scripts and custom shell scripts, following the AFNI-recommended processing pipeline^51^. The raw functional data from each scan were first subjected to slice-timing correction: the first 12 TRs of each scan were removed to exclude the unstable, noisy signals at the beginning of the acquisition, and the time series for each slice were then temporally shifted using the first slice as the reference. Subsequently, the data were corrected for head and body motion using a slice-wise realignment procedure that allowed for shifts and rotations within each slice, with the reference volume selected as the one containing the lowest number of outlier voxels, after which the motion-corrected functional data were aligned to the high-resolution anatomical image that had been intensity-uniformized. To further improve the signal-to-noise ratio of the functional data, the following preprocessing steps were applied^52^: [1] resampling the data to an isotropic voxel size of 1.5 mm×1.5 mm×1.5 mm; [2] applying a mask derived from the structural image; [3] applying Gaussian smoothing with a kernel radius of 2.0 mm; and [4] scaling the intensity values to a range between 0 and 200.

The general linear model (GLM) was used to estimate the response amplitude (β values) of each voxel to each block type (SELF, VF, NF, and scrambled) with motion parameters included as nuisance regressors and a MION-based hemodynamic response function (HRF) applied. To improve statistical power, the following preprocessing steps were implemented: [1] detrending using second-order polynomials; [2] removal of the first block of each scan; and [3] exclusion of any scan with head motion > 1 mm.

Individual anatomical volumes were registered to the NIMH macaque template (NMT v2) using the AFNI Animal Warper, and cortical masks were defined using the NMT cortical atlas in template space ^51^. For whole-brain statistical analysis, functional data were aligned to the MNI–Paxinos macaque template brain. Given the small sample size (N = 3), we focused on single-subject analyses using a Linear Mixed-Effects (LME) modeling approach^53^. We specified a model with two within-subject factors: face type (SELF, VF and NF blocks) and training stage (Stage 1, Stage 2, and Stage 3), including their interaction. Random effects were incorporated to account for inter-subject variability, comprising a random intercept and random slopes for stage across subjects. This yielded the following LME model equation: functional response ∼ face_type × stage + (1 + stage | subject). The level of significance was set at *P* < 0.01.

Based on the whole-brain analysis results, we focused on activity in SII/PI (the secondary somatosensory cortex and posterior insula complex), V2 (visual area 2), V4 (visual area 4), TE (anterior inferotemporal cortex), TEO (posterior inferotemporal cortex), and STS (superior temporal sulcus), for which a bilateral anatomical mask was defined to guide further signal analysis. The SII/PI mask was manually delineated on the macaque template (NMT v2) based on slices containing significant voxels, with the union of the two corresponding brain regions (SII and insula) in the template used as the anatomical reference. All remaining anatomical masks were delineated based on established landmarks in the NMT v2 template. Then, regions of interest (ROIs) were defined as the top 10% of voxels showing the strongest preference for self-face over scrambled images, based on stage-3 data within the anatomical masks described above. Statistical analyses were conducted on the relative response amplitudes of these voxels across scans for each ROI. The effect index for face X versus face Y was calculated as: [β(X) − β(Y)] / (abs[β(X)] + abs[β(Y)]). The face coefficient was computed as: β(face) − β(scrambled). Statistical significance was set at *P* < 0.05.

Dynamic causal modeling (DCM) analysis was performed using the DCM module in SPM12 software^54^. The data used were preprocessed with AFNI, the pipeline and parameters were consistent with those used previously. The voxel mask used for signal extraction was derived from voxels showing significant responses to any experimental stimulus (F contrast, p < 0.05). Because a contrast agent was administered to the subjects during the MRI scan, we used a custom-made MION kernel instead of the default HRF function in SPM. Using V1 as the input node for visual information, V2 and SII-PI as downstream brain regions, we constructed a three-node model and investigated the modulatory effects of the three face stimuli on the connections within the model. DCMs were estimated separately for each functional run, and all runs from the three monkeys were pooled into a single group-level analysis, so that the estimated modulatory effects reflect connectivity changes that are consistent across runs rather than a single scanning session. Model comparison and parameter inference were ultimately conducted using the Bayesian Model Averaging (BMA) method^55,56^.

