## supplement S1 S2 S3 TABLE S1 for "Cortical areas for self-recognition in the monkey brain"

**Figure S1**

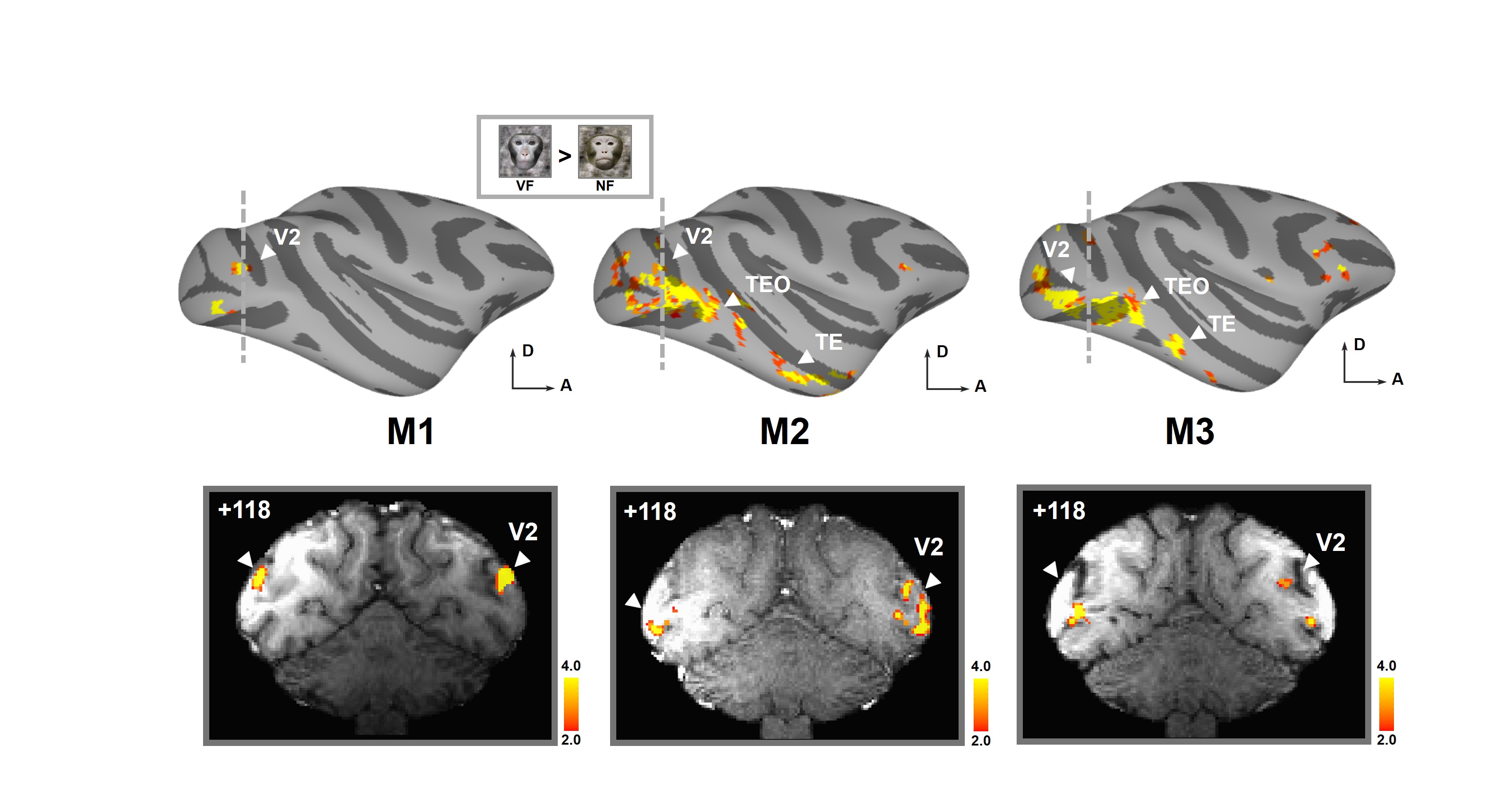

**Figure S1. Cortical activations in VF vs. NF contrast in MSR-capable monkeys.**

Inflated hemispheres showed the regions that responded significantly more to visual familiar (VF) face than to unfamiliar (non-familiar, NF) face using an LME model in the three monkeys with MSR ability (uncorrected, *P* < 0.01). The color bar denotes t-statistic values. V2, visual area 2; TE, anterior inferotemporal cortex; TEO, posterior inferotemporal cortex.

**Figure S2**

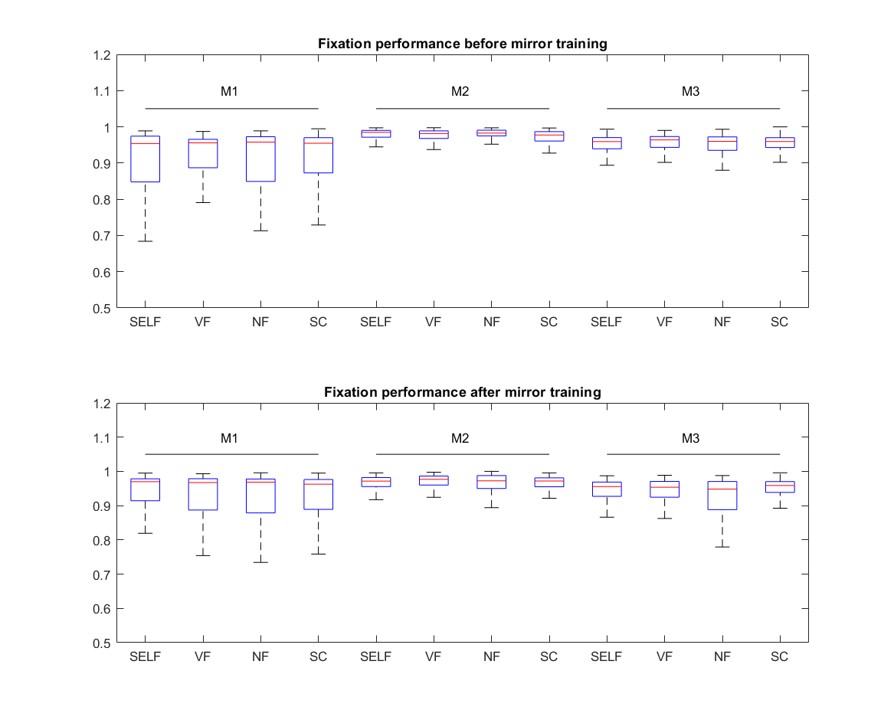

**Figure S2. Fixation performance across conditions in monkeys**

Fixation performance is expressed as the percentage of time within each 24-s stimulus block during which gaze remained within the fixation window. Saccades and fixations outside this window, as well as blinks, reduced this percentage. Data distributions are shown using boxes representing the interquartile range (with the top and bottom edges corresponding to the 75th and 25th percentiles, respectively) and horizontal lines indicating the sample median.

**Figure S3**

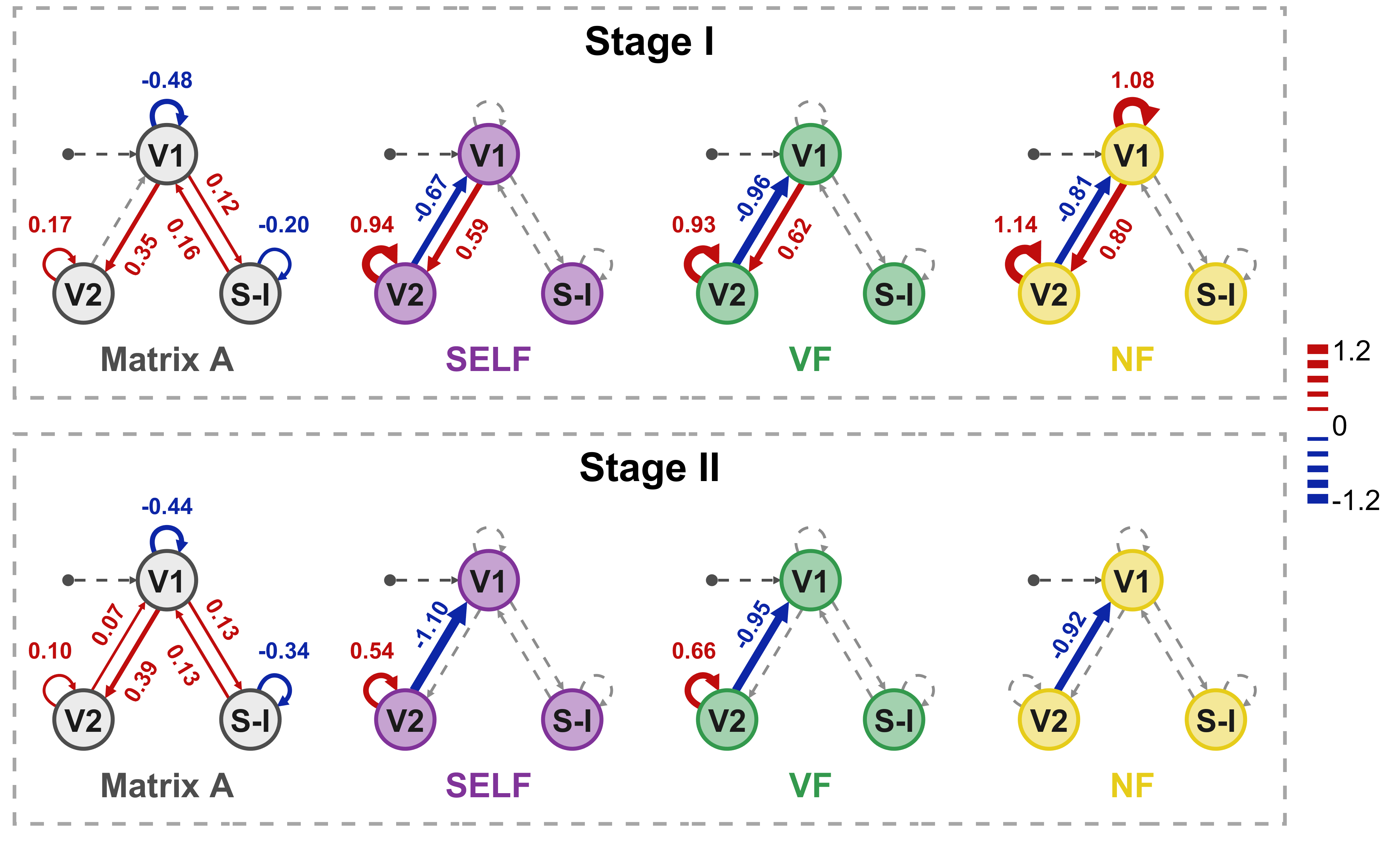

**Figure S3. Stage-specific modulatory effects of face stimuli at Stage I and Stage II**

For each stage, the intrinsic connectivity of the three-node model (V1, V2, and S-I) is shown as “Matrix A”, and the modulatory effects of SELF, VF, and NF are shown separately. Numbers are parameter estimates; colored arrows are shown for connections with posterior probability > 0.85, whereas gray dashed arrows indicate connections without significant modulation. The color bar denotes modulation strength (red, positive; blue, negative). S-I, SII-PI complex.

**Table S1.** Stereotactic coordinates of the peak voxel (SELF versus VF or NF) of SII-insula in the left and right hemispheres of the three monkeys tested. Coordinates are in NMT template space.

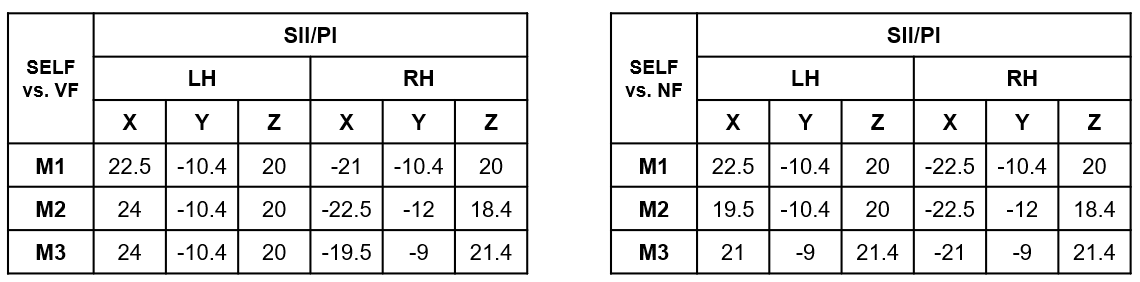

**Video S1.** Typical video showing the behavioral performances of a monkey during MSR training and subsequent mirror mark test.
